# In Vivo Antiviral Efficacy of LCTG-002, a Pooled, Purified Human Milk Secretory IgA product, Against SARS-CoV-2 in a Murine Model of COVID-19

**DOI:** 10.1101/2023.08.25.554813

**Authors:** Viraj Mane, Rikin Mehta, Nadine Alvarez, Vijeta Sharma, Steven Park, Alisa Fox, Claire DeCarlo, Xiaoqi Yang, David S. Perlin, Rebecca L.R. Powell

## Abstract

Immunoglobulin A (IgA) is the most abundant antibody (Ab) in human mucosal compartments including the respiratory tract, with the secretory form of IgA (sIgA) being dominant and uniquely stable in these environments. sIgA is naturally found in human milk, which could be considered a global resource for this biologic, justifying the development of human milk sIgA as a dedicated airway therapeutic for respiratory infections such as SARS-CoV-2. In the present study, methods were therefore developed to efficiently extract human milk sIgA from donors who were either immunologically naïve to SARS-CoV-2 (pooled as a control IgA) or had recovered from a PCR-confirmed SARS-CoV-2 infection that elicited high-titer anti-SARS-CoV-2 Spike sIgA Abs in their milk (pooled together to make LCTG-002). Mass spectrometry determined that proteins with a relative abundance of 1.0% or greater were all associated with sIgA. None of the proteins exhibited statistically significant differences between batches. Western blot demonstrated all batches consisted predominantly of sIgA. Compared to control IgA, LCTG-002 demonstrated significantly higher binding to Spike, and was also capable of blocking the Spike - ACE2 interaction in vitro with 6.3x greater potency compared to control IgA (58% inhibition at ∼240ug/mL). LCTG-002 was then tested in vivo for its capacity to reduce viral burden in the lungs of K18+hACE2 transgenic mice inoculated with SARS-CoV-2. LCTG-002 was demonstrated to significantly reduce SARS-CoV-2 titers in the lungs compared to control IgA when administered at either 250ug/day or 1 mg/day, as measured by TCID50, plaque forming units (PFU), and qRT-PCR, with a maximum reduction of 4.9 logs. This innovative study demonstrates that LCTG-002 is highly pure, efficacious, and well tolerated in vivo, supporting further development of milk-derived, polyclonal sIgA therapeutics against SARS-CoV-2 and other mucosal infections.

## 1. INTRODUCTION

### 1.1 Secretory IgA structure and function

Immunoglobulin A (IgA) is one of 5 human antibody (Ab) subclasses. Healthy humans naturally produce IgA in response to infections. Secretory IgA (sIgA) is the predominant Ab class found in human mucosal compartments (e.g. the respiratory tract, gastrointestinal and genitourinary tracts, and the oral/nasal cavity), and is distinguished from monomeric IgA by the scaffolding of IgA monomers into an end-to-end formation (typically dimeric) via an intermediary J-chain, which is wrapped in Secretory Component (SC) as it is secreted into mucosae. SC confers protection to sIgA, stabilizing it in relatively harsh mucosal environments that can otherwise degrade biomolecules via low pH, proteases, ciliary activity, and mucus entrapment [1]. This protective SC-conferred mucosal stability does not occur for IgG, which is the predominant Ab class found in the blood, or any other monomeric Ig class. sIgA primarily acts in the respiratory tract or other mucosae by recognizing pathogens (i.e., viruses, bacteria, fungi, and parasites) in an antigen-specific manner and neutralizing their activity, preventing their attachment to target cells through a process called immune exclusion [2]. The neutralization effect has been shown to be significantly more potent using sIgA compared to monomeric IgA, likely due in part to its enhanced durability and increased avidity as a polymer [3, 4]. Unique among human Ab classes, sIgA can also neutralize intracellular viruses by interfering with their replication and/or assembly [5]. sIgA also mediate various non-neutralizing functions via its Fc domain via interactions with Fc receptors (FcRs) [6–9]. The Fc region of sIgA is also a potent activator of the alternative pathway of complement [10]. The abundant glycans on sIgA also activate the lectin pathway of complement and non-specifically contribute to pathogen clearance [3, 4, 10].

Respiratory pathogens must access the nostrils and/or mouth to infect airway cells, replicate, and subsequently penetrate the nasal cavity, deeper airways, and lungs. Therefore, by (a) presenting an sIgA barrier and (b) physically entrapping pathogens and particles in mucus, the nasal cavity’s mucosal layer provides two key layers of early defense against respiratory infections.

### 1.2 IgA Deficiency

The importance of sIgA in respiratory infections is illustrated in patients with common variable immune deficiency (CVID), a rare, chronic, and incurable heterogeneous immune disorder driven by defective B-cell differentiation leading to deficient serum levels of IgA and IgG [11]. A subset of CVID patients additionally have an IgM deficit [12, 13]. Due to compromised Ab immunity, the most prominent clinical sequelae are respiratory infections, which afflict 73% of individuals with CVID [14]. Sino-pulmonary diseases such as pneumonia, chronic sinusitis, chronic otitis media, and bronchitis are the greatest contributors to patient morbidity and mortality, both of which are worse in children [15, 16]. Among respiratory pathogens, bacterial infections are most prevalent, followed by viral infections. Interestingly, low serum IgA levels are correlated with an increased risk of respiratory infections of viral origin [11]. CVID also leads to poor antibody response to vaccination [15]. The primary treatment for CVID is immunoglobulin replacement therapy (Ig-RT), which can be administered intravenously or subcutaneously [14]. Although it has been demonstrated that Ig-RT can achieve adequate serum IgG levels, as well as reduce the number of respiratory infections, hospitalizations, and antibiotic use, there are considerable shortcomings to current formulations of Ig-RT [11]. Of note, Ig-RT is not delivered to the airway, which is one of the most prevalent areas of infection in CVID patients. Furthermore, commercially available Ig preparations contain IgG, with little to no IgA.

Lastly, the IgA in human plasma, from which Ig used for Ig-RT is sourced, is primarily in the monomeric form and not the dimeric secretory form. For these reasons Ig-RT does not effectively control respiratory infections, and CVID patients on Ig-RT still experience significant mortality and serious morbidity (e.g., bronchiectasis) due to respiratory infections [14], underscoring the need for infection prevention in the nasal cavity. Another condition characterized by deficient IgA production and increased risk of respiratory infections is Selective IgA Deficiency (SIgAD). In the symptomatic SIgAD subset, frequent infections occur secondary to the IgA deficiency and are treated with extensive use of antibiotics and administration of intravenous or subcutaneous Ig replacement therapy [17].

### 1.3 COVID-19 and SARS-CoV-2-specific IgA

SARS-CoV-2, the causative agent of COVID-19, infects human airway cells via the receptor binding domain (RBD) of its Spike protein binding to human receptor ACE2 [18]. COVID-19 infection provokes mucosal immunity including the production of protective, SARS-CoV-2-specific IgA as confirmed in COVID-19 patient cohorts in multiple countries. In COVID-hospitalized patients in the UK, SARS-CoV-2-specific nasal IgA concentrations increased over baseline as early as 14 days after infection and remained elevated for up to 9 months following infection [19]. In a study of 338 triple-vaccinated Swedish health care workers, high levels of wild-type spike-specific mucosal IgA were correlated with protection against subsequent omicron breakthrough infection [20]. In the same vein, clonally expressed dimeric IgA from COVID-convalesced US patients were more effective at RBD binding and SARS-CoV-2 neutralization compared to monomeric IgA [21].

While these clinical studies highlight the importance of dimeric sIgA in protective responses to COVID-19, they also underscore the consequences of IgA deficiency in the context of CVID patients. Indeed, COVID-19 vaccines tested in a Spanish CVID cohort were found to be less efficient at inducing COVID Spike-specific serum Abs as compared to healthy controls [22] and a study in a Swiss CVID cohort found a similar outcome [23]. While these studies measured serum and not mucosal samples, they emphasize the overall deficiency in CVID patients’ generation of SARS-CoV-2-neutralizing antibody responses to vaccines and underscore the potential value of sIgA replacement for these patients.

### 1.4 Study Goals

sIgA is the predominant Ab class in human milk, at a concentration of up to 0.8 g/L in the first year of lactation [24], making human milk the only naturally abundant source of polyclonal human sIgA. The therapeutic potential of human milk sIgA against SARS-CoV-2 was previously tested by demonstrating that milk from COVID-recovered donors contained IgA specific for the receptor-binding domain (RBD) of SARS-CoV-2 Spike protein, and the IgA was in the secretory form [25]. Furthermore, human milk samples contain persistent Spike-specific sIgA titers for 12 months or more after infection [26, 27]. Therefore, a medical preparation of polyclonal human sIgA, delivered directly to the nasal cavity, is a promising avenue for pre- and post-exposure prophylaxis against COVID-19.

The present study sought to characterize LCTG-002, a therapeutic candidate consisting of purified polyclonal sIgA extracted from pooled human milk according to a proprietary, validated and patented production method. This milk was obtained from lactating people previously infected with SARS-CoV-2, but not vaccinated against COVID-19, who had been shown to exhibit high titers of Spike-specific milk sIgA [26]. LCTG-002 and control sIgA batches were characterized for batch production consistency, purity, and stability, then assessed in vitro to demonstrate binding to SARS-CoV-2 Spike protein and neutralization of Spike binding to ACE2 receptor. Finally, LCTG-002 was delivered to the airways of ACE2 transgenic mice before and after inoculation with wild type SARS-CoV-2 to ascertain in vivo antiviral efficacy in a prophylactic dosing regimen.

## 2. MATERIALS AND METHODS

### 2.1 Selection of human milk donors and milk procurement

Milk samples used for this study were obtained from study participants who either (1) had a SARS-CoV-2 infection confirmed by an FDA-approved COVID-19 PCR test 3-8 weeks prior to the initial milk sample collection, or (2) had no history of SARS-CoV-2 infection (control samples) as described previously [26].

### 2.2 LCTG-002 preparation

LCTG-002 was prepared by purifying IgA from pooled milk samples based on methods disclosed in patent US 11,124,560, dialyzed into PBS pH. 7.0, then 0.2 μm filtered.

### 2.3 LCTG-002 protein imaging

A normalized 2ug sample of LCTG-002 was mixed with an equivalent volume of 2X laemmli loading buffer containing BME. 1ug LCTG-002 samples, as well as a 8uL sample of Precision Plus Protein Unstained Standard by BioRad, were loaded onto a Mini-PROTEAN TGX Stain-Free Gel by BioRad and run with 1X Tris/Glycine SDS Buffer at 200 Volts for 35 minutes. The gel was stained with GelCode Blue Safe Protein Stain for 1h at room temperature with gentle mixing, the stain was poured off, the gel was unstained with distilled water overnight at room temperature with gentle mixing, and gel was imaged with the Coomassie-stained program from Syngene’s G BoxMini. The image was analyzed with ImageJ to determine relative purity as follows: image was cropped to remove any white background regions, and background was subtracted starting with a rolling ball radius of 100 and adjusted as needed for lightness/darkness. Gel lanes of interest were selected and the peak regions were selected and separated from other peaks, while keeping note of major peaks predicted to be near the corresponding molecular weight to the target protein. The area of the chosen peaks were determined via ImageJ software and the data were transferred onto a spreadsheet to calculate relative purity percentage: ((Sum of peaks of interest)/(sum of all peaks))*100.

For the native gel western blot analysis, 0.5ug for the IgA blot and 5ug for the SC blot of LCTG-002 was loaded on a 7.5% pre-casted Gel Mini Protein gel TGX (BioRad) in native gel buffer. Gels were transferred by iBlot2 to a PVDF membrane and blocked with 5% nonfat milk for 1h. Blotting was performed using 1:10000 anti-IgA HRP (EMD Milipore) or 1:3000 anti-SC HRP (Nordic Mubio). A standard ECL substrate was used to develop (Biorad) and blot was visualized on a BioRad Imaging System.

### 2.4 Mass spectrometry

Samples were analyzed as described previously [28]. In brief, 10ug samples of LCTG-002 (3 replicates per group) were run ∼1cm into an SDS-PAGE. Gels were stained with Coomassie blue and gel pieces excised. Proteins were digested with in-gel trypsin and resulting peptides excised and analyzed on a Thermo Scientific Eclipse mass spectrometer. Data were analyzed using Maxquant and Perseus to provide an FDR-controlled method to compare differences in protein expression between the two groups. Absolute protein amounts were determined using the label-free iBAQ component of Maxquant.

### 2.5 Spike binding assays

ELISA was performed to measure Spike binding as previously described [26]. An ELISA-based proxy neutralization assay that detects Abs specific for the Spike RBD that can prevent RBD-ACE2 binding was also performed, using serially diluted purified LCTG-002, following manufacturer’s protocol (Genscript). Plates were read by Powerwave plate reader, and the percent inhibition was calculated according to the positive and negative controls from the kit. LCTG-002 binding to SARS-CoV-2 Spike protein was also measured via optimized Luminex assay as described [27].

### 2.6 Mouse studies

The K18+hACE2 mouse model (Jackson Laboratory) was chosen based on internal data and literature indicating that SARS-CoV-2 productively replicates in this mouse model expressing human ACE2 [29]. All animal experiments were performed under Hackensack Meridian Health Center for Discovery and Innovation (CDI) IACUC approval and the animals were housed at the CDI Research Animal Facility which is accredited by the Association for Assessment and Accreditation of Laboratory Animal Care International (AAALACi). All mice were housed in individual ventilated caging (IVC) units at CDI and handled under sterile conditions for a minimum of 72 hours for acclimation prior to any study start. For necropsy, mice were humanely euthanized by cervical dislocation and nasal washes were collected. Lungs were aseptically collected and weighed. Half of the lungs were placed in 2ml screw cap tubes and stored at −80□. The other half were weighed, placed in a GentleMACS Octo dissociator tube containing 2.5ml of DMEM + 2% FBS +1x AB/AM and homogenized (modified from [30]). 0.475ml was transferred to a screw cap tube and 0.025 ml of Proteinase added then heat inactivated for at least 60min at 65C for qRT-PCR. 0.5ml of untreated supernatant was collected for PFU and TCID50 assays. 0.1ml of the nasal washes were used for these assays, which were described previously [31].

#### 2.6.1 SARS-CoV-2 inoculum

SARS-CoV-2 virus strain USA-WA1/2020 (P4, Catalog#: NR-53873, Lot#:70039812, 9.2X105 TCID50/ml (Vero E6) Lot#:70039812) was obtained from BEI Resources. SARS-CoV-2 was gently thawed on ice and used to directly inoculate mice. Mice were anesthetized by inhalation of vaporized isoflurane and intranasally infected with 50μl of the virus stock to a final infection dose of 4.6 × 10^4^ TCID50.

#### 2.6.2 Administration of test materials to mice

On each day of dosing, LCTG-002 was thawed to room temp and 0.05ml of LCTG-002 or PBS control was dosed into the mouse nares (0.025ml per naris) intranasally in the morning and 0.05ml intratracheally in the evening as outlined in Table 1. Mice were anesthetized by inhalation of vaporized isoflurane for LCTG-002 administration. For Nirmatrelvir administration, a 300mg pill was pulverized using a sterile mortar and pestle then suspended in sterile water. Nirmatrelvir suspension was orally administered at 250mg/kg in 10ml/kg using a tuberculin syringe and a 20g popper feeding needle.

**Table 1.**
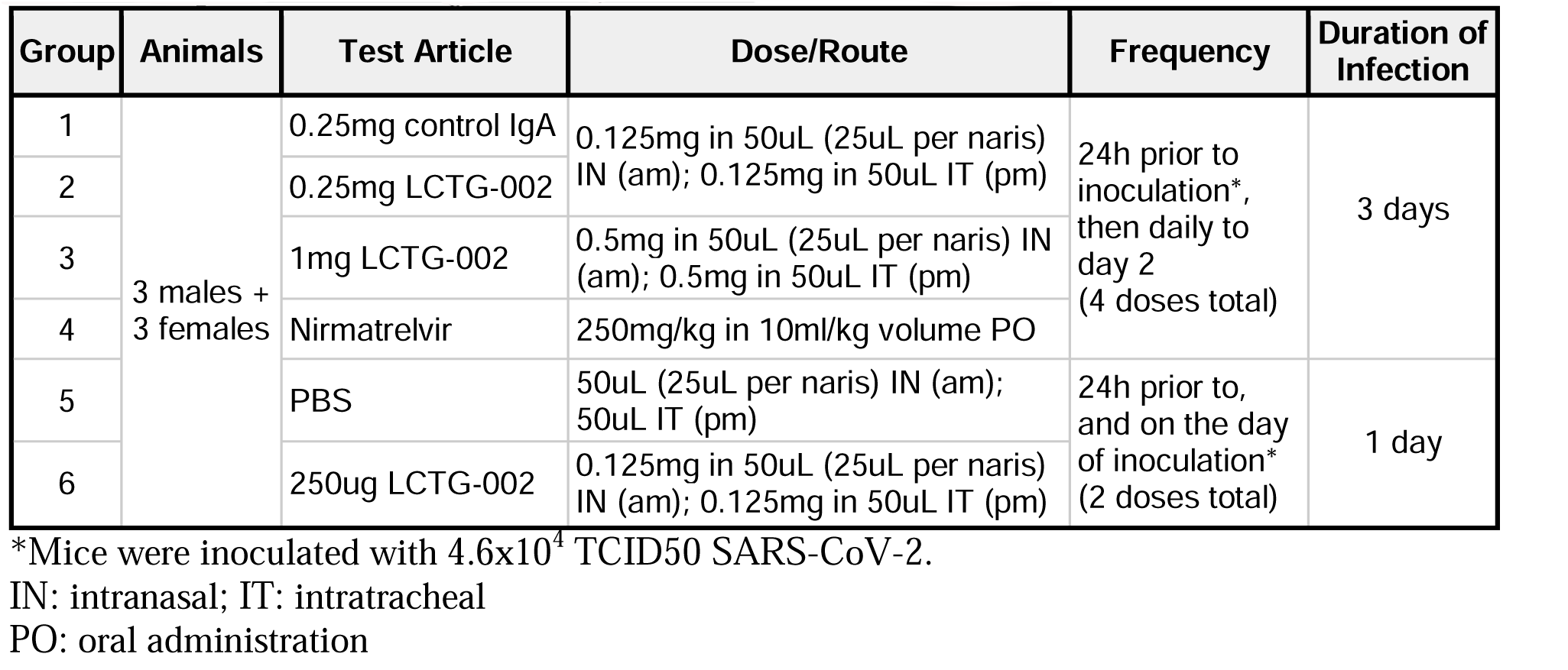
Experimental design for K18+hACE2 mouse studies.

## 3. RESULTS

### 3.1 Characterization of pooled, purified milk IgA

IgA was extracted from pooled human milk samples from either pre-pandemic, COVID-naive lactating people as a negative control (control IgA) or COVID-recovered lactating people (LCTG-002) based on methods disclosed in patent US 11,124,560. Samples were run on SDS-PAGE, stained with Coomassie blue, and imaged. A pattern of 3 protein bands was consistently visible in both batches (Fig. 1) corresponding from top to bottom to the known molecular weights of SC, heavy alpha chain, and light chain. Two additional batches were prepared from pooled human milk, imaged, and demonstrated identical protein bands (data not shown). Protein purity was assessed across all 4 batches by SDS-PAGE imaging and ranged from 93.2-99.2% across batches (data not shown). Samples from the control IgA and LCTG-002 batches were also evaluated by western blot to compare sIgA content. The blot was immunostained for human IgA and the signal was identical in both batches (Fig. 2, left panel). A separate blot was immunostained for human SC and the signal was also identical in both batches (Fig. 2, right panel), confirming that both batches had similar human IgA content which was predominantly in the secretory (sIgA) form. The upper smear in the left panel is indicative of differently-glycosylated sIgA isoforms. The lower band above 130 kDa likely represents IgA monomers (left panel); monomers are not bound by secretory component, which explains the absence of a corresponding band in the right panel.

**Figure 1.**
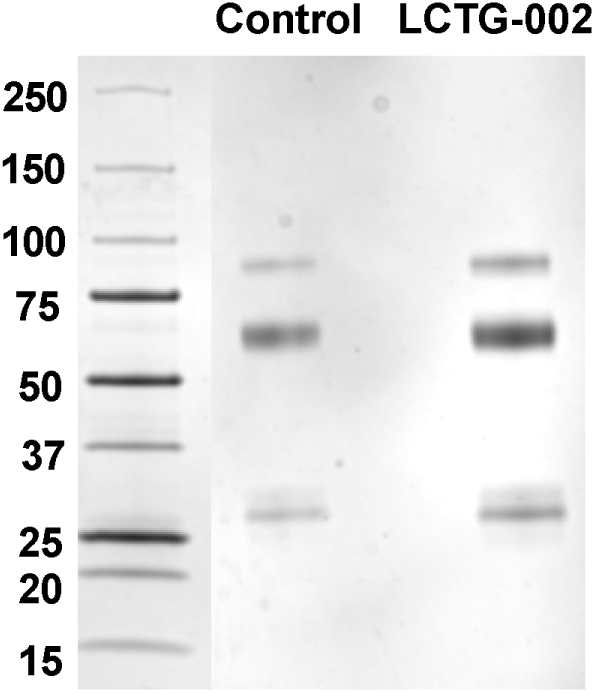
Imaging and purity assessment of human antibody extracts. sIgA was purified from pooled human milk samples from donors that were either COVID-naive (Control IgA) or COVID-recovered (LCTG-002), aliquots were run on SDS-PAGE, then stained with Coomassie blue to visualize sIgA protein bands (right lanes) relative to a protein marker (left lane).

**Figure 2.**
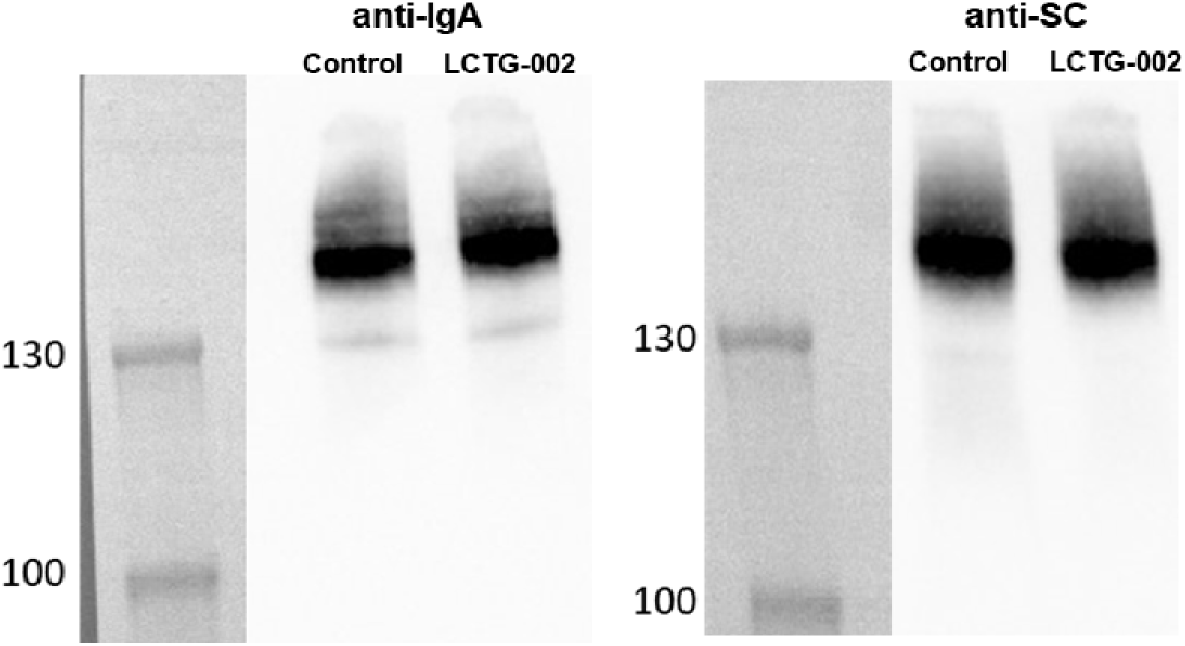
Western blot analysis of human milk-derived IgA. Aliquots of pooled, purified IgA from human milk obtained from COVID-naive (Control) or COVID-recovered (LCTG-002) individuals were assessed by native gel western blots on a 7.5% pre-casted Gel Mini Protein gel TGX (BioRad) in native gel buffer. Gels were transferred by iBlot2 to a PVDF membrane and blocked with 5% nonfat milk for 1h. Blotting was performed using 1:10000 anti-IgA HRP (EMD Milipore) or 1:3000 anti-SC HRP (Nordic Mubio). A standard ECL substrate was used to develop (Biorad) and blot was visualized on a BioRad Imaging System.

To quantitatively assess relative protein abundance, mass spectrometry was performed on the control IgA and LCTG-002 batches and data were analyzed using the search engine MaxQuant with label-free quantification to generate iBAQ values. Proteins with a relative abundance of 1.0% or greater were plotted in descending order. These proteins were all associated with sIgA, specifically: immunoglobulins, SC (which is a cleavage product of Polymeric immunoglobulin receptor (PIGR)), and J chain (JCHAIN) (Fig. 3, Panel A). None of the proteins exhibited statistically significant differences between batches, establishing that protein composition was comparable.

**Figure 3.**
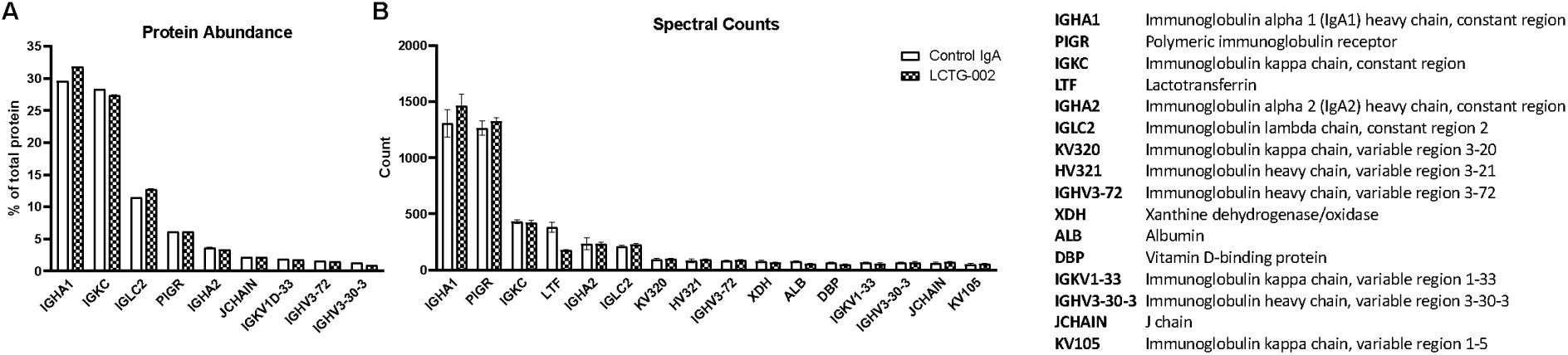
Mass spectrometry characterization of antibody preparations. Relative protein abundance in Control IgA and LCTG-002 batches was calculated from mass spectrometry data by dividing each protein’s individual iBAQ value by the total iBAQ sample value. An abundance of 1.0% was used as the lower threshold. Averages and standard deviations were calculated from triplicate samples and plotted (Panel A); note the error bars were plotted but are too small to be visible at this scale. Peptides from Control IgA and LCTG-002 batches were also plotted in order of descending spectral count (Y-axis) with a spectral count of 50 used as an arbitrary lower threshold; averages and standard deviations were calculated from triplicate samples and plotted (Panel B).

An additional mass spec-based assessment of individual protein composition between control IgA and LCTG-002 batches was performed by using X!Tandem semi-quantitative peptide analysis of protein spectral counts. These are plotted in descending order (Fig. 3, Panel B). 12 out of 16 proteins were associated with sIgA and these proteins demonstrated similar spectral counts between batches.

Taken together, milk IgA was consistent across control IgA and LCTG-002 batches despite differing in various categories: (1) number of pooled donors per batch, (2) date of milk collection, (3) identity of the donors, (4) COVID-19 infection status, and (5) date of processing. This establishes that the purification methods are reproducible and the batches are highly pure and consistent with respect to protein composition.

### 3.2 In vitro assessment of pooled, purified milk IgA function

The binding specificity of LCTG-002 against SARS-CoV-2 Spike was assessed by ELISA. Using a secondary Ab specific for human IgA, significantly greater Spike-specific IgA binding was found for LCTG-002 (OD_450_ = 3.36 +/− 0.07) compared to control IgA (1.89 +/− 0.15; p = 0.0088), using 5ug of Spike per well (Fig. 4, Panel A).

**Figure 4.**
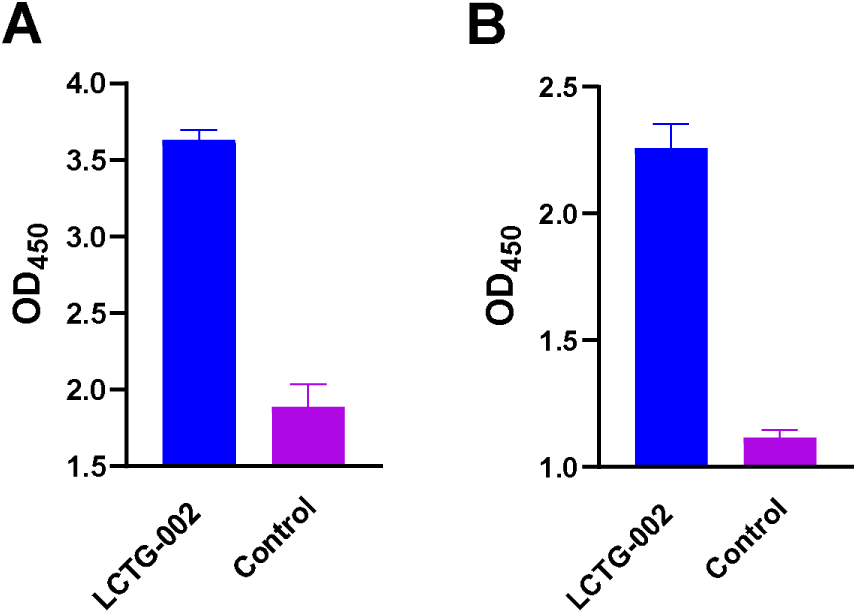
LCTG-002 binding to SARS-CoV-2 Spike protein. LCTG-002 (blue bars) or Control IgA (purple bars) were tested at 100 ug/mL for binding to Spike protein via ELISA with secondary antibody detecting human IgA (A) or human Secretory Component (B) which were quantified by OD450 (Y-axis). Groups were tested in triplicate, error bars denote standard error of the mean (SEM); p value between LCTG-002 and Control IgA is < 0.01 in both panels.

Using a secondary Ab specific for human SC to confirm that the Spike-specific human IgA were largely in secretory form (sIgA), it was similarly found that LCTG-002 exhibited significantly higher Spike-specific binding (2.26 +/−0.09) compared to control IgA (1.12 +/− 0.03; p = 0.0076) (Fig. 4, Panel B). The similarity between Panels A and B confirms that the pooled IgA are primarily in secretory (sIgA) form, consistent with our previous study [26]. For both IgA- and SC-specific ELISAs, control IgA and LCTG-002 were serially diluted and demonstrated a similar differential in binding across the tested concentrations (data not shown).

To further probe the biological characteristics of LCTG-002, an ELISA-based proxy neutralization assay was used to measure LCTG-002’s ability to compete with the interaction between ACE2 receptor and the Spike RBD. LCTG-002 demonstrated a maximum inhibition of 58% +/− 1.2% at 242ug/mL while control IgA did not exceed 6.5% +/− 0.6% inhibition (Fig. 5). Area under the curve analysis of inhibition curves found LCTG-002 activity to be 6.3x greater compared to control, further confirming the presence of Spike-specific neutralizing sIgA in LCTG-002.

**Figure 5.**
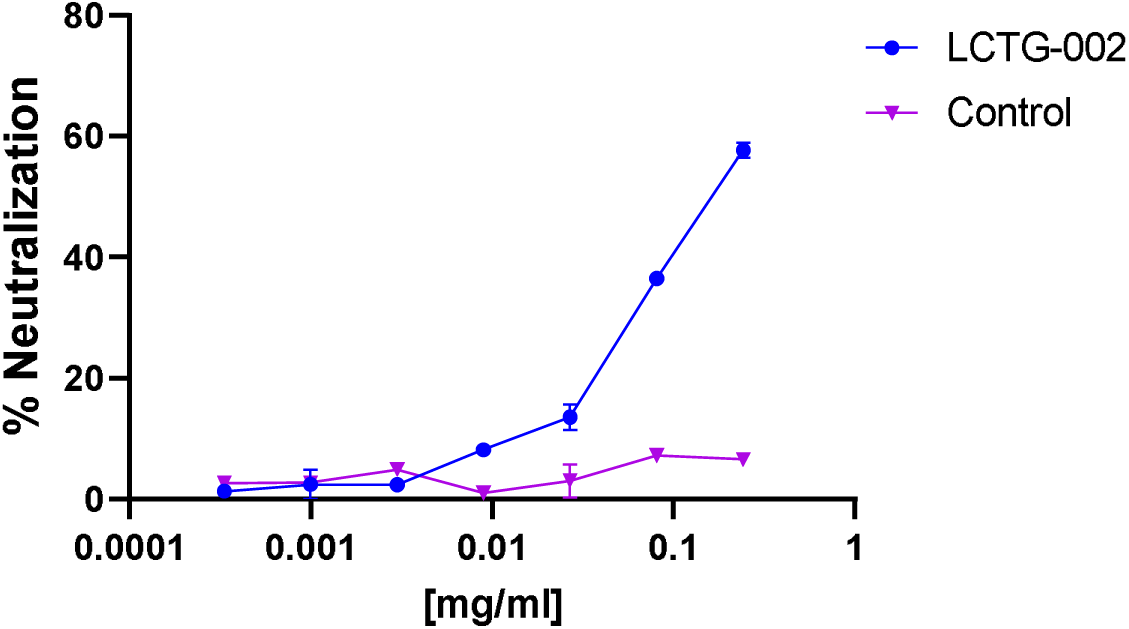
LCTG-002 inhibition of Spike-RBD interaction with hACE2. LCTG-002 (blue circles) or Control IgA (purple triangles) were tested at a range of dilutions (X-axis) for their ability to compete with SARS-CoV-2 Spike protein RBD interaction with hACE2 receptor (Y-axis). Groups were tested in triplicate, error bars denote SEM; p value between groups is < 0.001 at the highest concentration.

Taken together, these data demonstrate that LCTG-002 extracted from COVID-19-recovered milk donors is predominantly in secretory form (sIgA), exhibits significantly greater SARS-CoV-2 Spike binding specificity as compared to control IgA, and blocks the SARS-CoV-2 Spike RBD from binding its human ACE2 receptor target.

### 3.3 In vitro assessment of pooled, purified milk IgA stability

A Luminex xMAP bead-based assay was used to further compare the ability of LCTG-002 and control IgA to bind SARS-CoV-2 Spike. Frozen control IgA or LCTG-002 samples underwent 1 of 3 conditions prior to bead incubation: (1) thawing and direct incubation; (2) thawing followed by one freeze-thaw cycle prior to incubation; or (3) thawing followed by two freeze-thaw cycles prior to incubation. Control IgA or LCTG-002 was serially diluted from 100ug/mL to 0.137ug/mL to generate separate binding curves measuring IgA and secretory Ab activity (Fig. 6). These curves were virtually identical regardless of freeze-thaw history, demonstrating that LCTG-002 binding to Spike is highly stable and unaffected by multiple freeze-thaw cycles, ensuring that potential differences in freeze-thaw status between LCTG-002 and control sIgA batches (e.g. during preparation and/or transport) did not contribute to observed differences in activity in the studies described herein.

**Figure 6.**
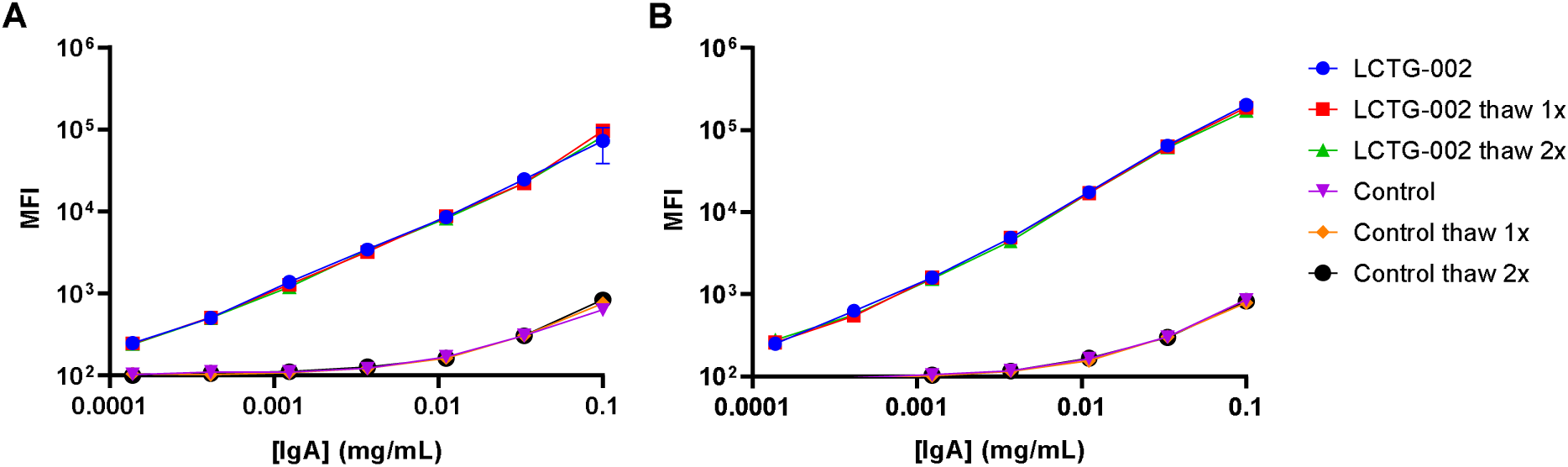
LCTG-002 stability and preservation of antibody binding. Control IgA and LCTG-002 were diluted into a range of concentrations (X-axis) and incubated directly with SARS-CoV-2 Spike-coated Luminex beads, or underwent one round of freeze-thaw (“thaw 1x”) or two rounds of freeze-thaw (“thaw 2x”) prior to bead incubation. Secondary antibodies were specific for human IgA (A) or human Secretory Component (B) with mean fluorescence intensity (MFI) plotted in log scale on the Y-axis. Groups were tested in triplicate; error bars denote SEM.

### 3.4 In vivo assessment of pooled, purified milk IgA efficacy against SARS-CoV-2

In order to test the potential antiviral efficacy of LCTG-002 against SARS-CoV-2 infection, K18+hACE2 transgenic mice were selected for use, based on literature indicating that SARS-CoV-2 productively replicates in this mouse model that expresses human ACE2 [29]. All mice were bright, alert, and responsive to manual restraint throughout the study, and no significant weight losses were observed in any group (data not shown).

PBS, control IgA, or LCTG-002 were administered both intranasally and intratracheally to model IgA deposition across the nasal cavity, airways, and lungs, which are targets of SARS-CoV-2 infection in humans. Administration occurred 1 day before, the day of, and each day after intranasal inoculation with an infectious dose of 4.6 × 10^4^ TCID50 SARS-CoV-2, as outlined in Table 1. A positive control group of mice received Nirmatrelvir, the SARS-CoV-2 protease inhibitor component of Paxlovid, by the oral route.

Viral titers were measured at the end of the study in the lungs 72 hours post inoculation via TCID50, plaque forming units (PFU), and qRT-PCR assay. Mice receiving 250ug/day of LCTG-002 exhibited statistically significant reductions in viral titer as measured by all 3 assays compared to mice receiving 250ug/day of control IgA (p=0.02 – p<0.0001; Fig. 7, Panel A).

**Figure 7.**
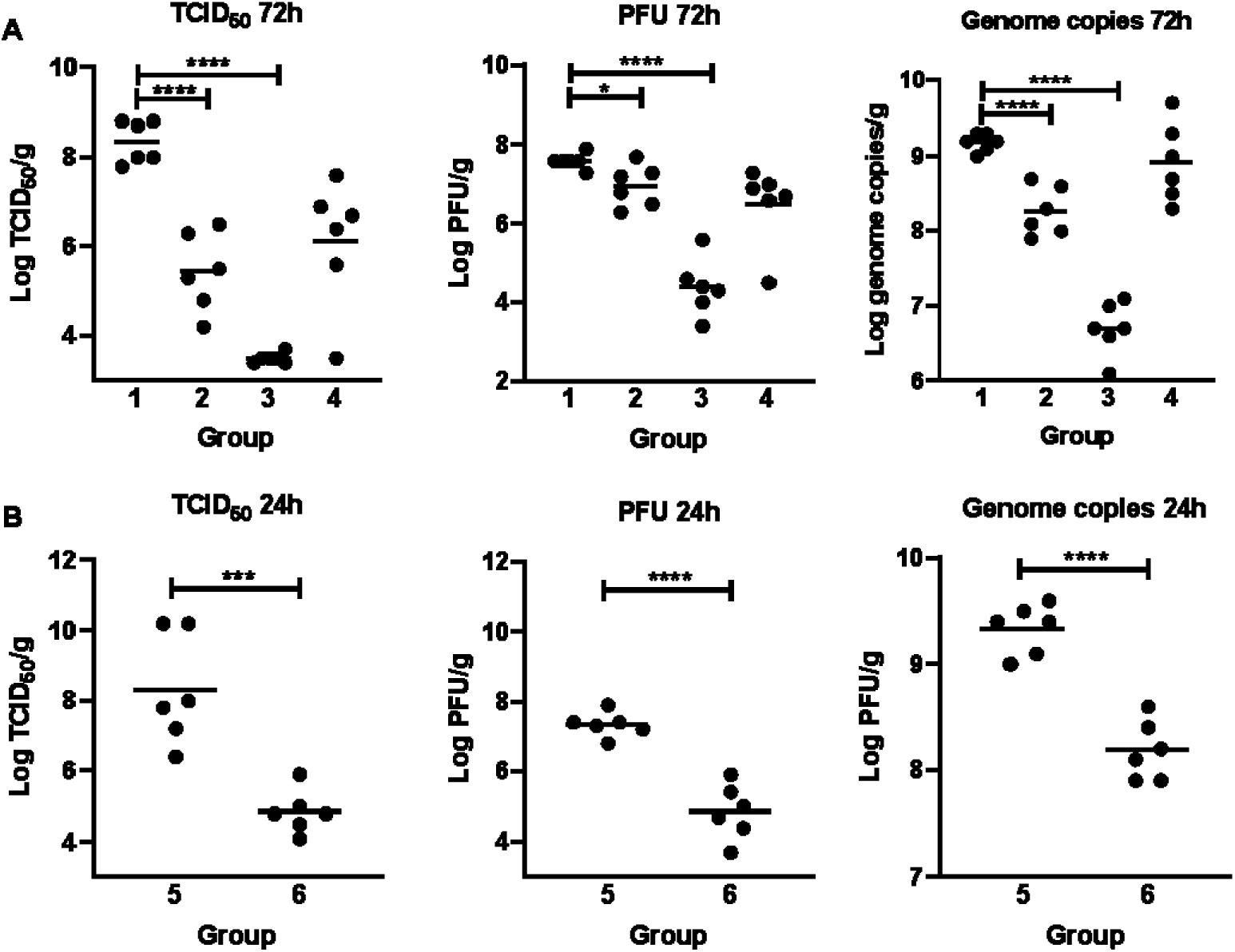
In vivo antiviral efficacy of LCTG-002 against SARS-CoV-2. Mouse lungs were collected and homogenized 72 hours (A) or 24 hours (B) after inoculation with 4.6 × 10^4^ TCID50 SARS-CoV-2, following a regimen of PBS, Control IgA, LCTG-002, or Nirmatrelvir (see Table 1) to measure viral burden via TCID50, PFU, and qRT-PCR assays. Viral titer is plotted in log scale; *P* values are denoted by asterisks as follows: *<0.05, **<0.01, ***<0.001, ****<0.0001.

Mice receiving 1mg/day of LCTG-002 exhibited even greater reductions in viral titer across all 3 assays, (P<0.0001), notably exhibiting a 4.9-log reduction in TCID50. Nirmatrelvir caused significant reductions in viral titer by TCID50 and PFU assays but not qRT-PCR, as similarly observed in prior unpublished studies using the same dose and route.

An additional mouse study was performed measuring viral titers in nasal washes and lung homogenates 24h after SARS-CoV-2 inoculation to determine whether prophylactic administration of LCTG-002 inhibited nasal and/or lung replication of SARS-CoV-2 in an earlier stage of infection. Compared to mice receiving PBS, mice receiving 250ug/day LCTG-002 exhibited statistically significant reductions in viral titers in lung across TCID50, PFU, and qRT-PCR assays (p=0.005 – p<0.0001; Fig. 7, Panel B). However, no significant reductions were observed in the nasal washes of mice receiving 250ug/day LCTG-002 compared to PBS treated mice (data not shown).

## 4. DISCUSSION

### 4.1 Summary of the data presented

LCTG-002, consisting of sIgA extracted from pooled human milk donated by lactating people that had recovered from COVID-19, was tested for activity against SARS-CoV-2 using in vitro and in vivo assays. The data presented herein support our hypothesis that pooled, polyclonal human milk sIgA is capable of both recognizing and neutralizing SARS-CoV-2. LCTG-002 exhibits specificity for SARS-CoV-2 Spike (Figs 4-6), consistent with the aforementioned studies in COVID-19 patient cohorts in the UK [19], Sweden [20], and the US [21]. This also establishes a proof of concept for the activity of pooled human milk sIgA against a multitude of other mucosal pathogens, as confirmed through our unpublished studies testing milk sIgA activity against various additional respiratory and enteric pathogens. The polyclonal, pooled nature of human sIgA in LCTG-002 is a critical aspect of translational relevance because polyclonal Ab comprises activity against a broad array of epitopes from numerous pathogens compared to monoclonal Ab. Conceptually, this should also result in a greater likelihood of cross-reactivity against emerging variants.

Of great significance in the present study is our landmark in vivo therapeutic efficacy testing of LCTG-002 against SARS-CoV-2 infection. LCTG-002 administered daily to K18+hACE2 transgenic mice both intranasally and intratracheally to model deposition across the respiratory tract prior to and after SARS-CoV-2 infection demonstrated significant antiviral efficacy (Fig. 7). LCTG-002 dosed at 250ug/day was effective in the lungs at two time points (24h and 72h post inoculation), suggesting it is capable of durable suppression of SARS-CoV-2 replication (Figure 7, panels A and B). As expected, the higher dose (1 mg/day) was more effective than the lower dose at suppressing lung viral titers (Fig. 7).

### 4.2 Implications of the data for further development of LCTG-002

IgA purity and composition were consistent across control IgA and LCTG-002 batches despite various differences in donors, dates of collection, COVID-19 infection status, etc. This underscores that while each donor and each milk sample are immunologically unique, the general parameters of purified milk Ab preparations such as concentration and composition are similar across donors as measured in Figs 1-3. As expected from a previous study [26], the IgA was predominantly in secretory (sIgA) form, as determined in Figures 2, 4, and 6. The consistency in IgA purity across batches ensures that analysis of LCTG-002 activity was not impacted by contaminants, and that activity against SARS-CoV-2 was driven by IgA and not other immunomodulatory human milk components. The Luminex assay (Fig. 6) demonstrated preservation of IgA activity throughout multiple freeze-thaw cycles. While this is a promising preliminary assessment of LCTG-002 stability especially in the absence of thermostabilizers and excipients, further formulation development must be performed and stability testing will be required.

### 4.3 Significance of the data in the context of other COVID-19 biologics

Convalescent plasma containing polyclonal Abs from COVID-19-recovered donors was employed as a therapeutic approach for patients hospitalized with COVID-19, particularly during the pre-vaccine pandemic period; notably, plasma contains mostly monomeric IgG. Convalescent plasma treatment was granted Emergency Use Authorization (EUA) by FDA on August 23, 2020. However, convalescent plasma from patients infected with SARS-CoV-2 from early in the pandemic was not capable of neutralizing COVID variant B.1.1.7 [32]. For this reason the EUA was revised to “limit authorization to the use of COVID-19 convalescent plasma with high titers of anti-SARS-CoV-2 antibodies for the treatment of COVID-19 in patients with immunosuppressive disease or receiving immunosuppressive treatment in either the outpatient or inpatient setting” [33]. However, the clinical benefit of convalescent plasma in this immunosuppressed cohort remains inconclusive, and systemic infusion therapy of primarily IgG may continue to demonstrate only limited efficacy against a mucosal infection such as COVID-19. Additionally, it is evident that unlike milk sIgA, specific Ab titers in blood against SARS-CoV-2 wane rapidly over time, limiting the potential donor pool for convalescent plasma in general [34].

A series of injected or infused COVID-19 monoclonal Ab products (bamlanivimab plus etesevimab, casirivimab plus imdevimab, sotrovimab, bebtelovimab, and tixagevimab plus cilgavimab (Evusheld)) have also been developed against earlier strains of SARS-CoV-2 but have had their Emergency Use Authorizations (EUA) withdrawn because they were not expected to provide significant protection against emerging Omicron variants [35]. This highlights the inherent limitation of recombinant monoclonal Ab development programs and underscores the clinical potential of an immuno-adaptive source of evolving Ab such as human milk.

### 4.4 Conclusions and translational relevance

This study establishes a promising proof-of-concept for LCTG-002 efficacy as a pre- and post-exposure prophylactic treatment for COVID-19, yet it highlights translational limitations. The ratio of infectious particles of SARS-CoV-2 per animal used in this study (4.6 × 10^4^ TCID50 per mouse) does not translate to the minimum infectious dose per human, which may number only in the hundreds of viral particles [36]. This is to be expected, given that mice are not the primary host for SARS-CoV-2 and a relatively high inoculum is therefore needed to model viral replication and pathogenesis. Consequently, LCTG-002 doses administered to mice in this study are not reflective of anticipated human dosing. Furthermore, clearance and biodistribution of airway-delivered exogenous polyclonal Ab in mice, which are obligate nose breathers, are not strongly representative of clearance in humans. Nevertheless, it is encouraging that daily administration of LCTG-002 to mice via intranasal and intratracheal routes was highly efficacious and very well tolerated, and future animal studies will assess LCTG-002 efficacy using various dosing and route regimens.

As Spike-specific sIgA titers are detectable in human milk for 12 months or more after SARS-CoV-2 infection [26, 27], with COVID-19 vaccination prior to or following infection boosting this response (Powell, et. al,., unpublished), human milk represents a sustainable and durable source of protective Ab against COVID-19 with an enormous potential donor pool globally. Unlike monomeric Ab, sIgA is highly stable and resistant to degradation in all relatively harsh mucosal compartments and fluids [37, 38]. This study establishes a novel proof of concept for airway administration of polyclonal, pooled sIgA from human milk as a well-tolerated and repeatable prophylactic antiviral treatment, with broader applicability beyond SARS-CoV-2.

## ACKNOWLEDGEMENTS

We acknowledge the support of the Bioexpression and Fermentation Facility (BFF) in the Department of Biochemistry and Molecular Biology at the University of Georgia for protein purification and characterization. We acknowledge the support of the Biological Mass Spectrometry Facility at the Center for Advanced Biotechnology and Medicine at Rutgers, The State University of New Jersey, for mass spectrometry of IgA samples. Lastly, we acknowledge the following personnel at Hackensack Meridian Health Center for Discovery and Innovation for their contributions to mouse efficacy studies: Enriko Dolgov MD, Alberto Rojas-Triana MS, Camila Mendez Romeo DVM, Taylor Tillery BS, and Kira Goldgirsh MS. As always, we thank our study participants for their generous milk donations.

